# Real-time DNA barcoding in a remote rainforest using nanopore sequencing

**DOI:** 10.1101/189159

**Authors:** Aaron Pomerantz, Nicolás Peñafiel, Alejandro Arteaga, Lucas Bustamante, Frank Pichardo, Luis A. Coloma, César L. Barrio-Amorós, David Salazar-Valenzuela, Stefan Prost

## Abstract

Advancements in portable scientific instruments provide promising avenues to expedite field work in order to understand the diverse array of organisms that inhabit our planet. Here we tested the feasibility for *in situ* molecular analyses of endemic fauna using a portable laboratory fitting within a single backpack, in one of the world’s most imperiled biodiversity hotspots: the Ecuadorian Chocó rainforest. We utilized portable equipment, including the MinION DNA sequencer (Oxford Nanopore Technologies) and miniPCR (miniPCR), to perform DNA extraction, PCR amplification and real-time DNA barcode sequencing of reptile specimens in the field. We demonstrate that nanopore sequencing can be implemented in a remote tropical forest to quickly and accurately identify species using DNA barcoding, as we generated consensus sequences for species resolution with an accuracy of >99% in less than 24 hours after collecting specimens. In addition, we generated sequence information at Universidad Tecnológica Indoamérica in Quito for the recently re-discovered Jambato toad *Atelopus ignescens*, which was thought to be extinct for 28 years, a rare species of blind snake *Trilepida guayaquilensis*, and two undescribed species of *Dipsas* snakes. In this study we establish how mobile laboratories and nanopore sequencing can help to accelerate species identification in remote areas (especially for species that are difficult to diagnose based on characters of external morphology), be applied to local research facilities in developing countries, and rapidly generate information for species that are rare, endangered and undescribed, which can potentially aid in conservation efforts.

## Introduction

Biodiversity is defined as the variety of life found on Earth, including variation in genes, species, and ecosystems. While about 1.9 million species have been described to date, there are an estimated 5-30 million species in total on the planet, with most of the diversity contained within tropical rainforests [1], [2], [3]. For instance Ecuador, despite its small size of 283,561 km^2^ (roughly 1.5% of South America), is one of the most biologically diverse countries in the world [4], [5]. Biodiversity is fundamentally important to natural and agro-ecosystems; it provides humans with an array of foods and materials, contributes to medical discoveries, furnishes the economy, and supports ecological services that make life on our planet possible [6]. Today species are going extinct at an accelerated rate because of environmental changes caused by human activities including habitat loss, spread of non-native species, pollution, and climate change [7], [8]. All of these threats have put a serious strain on the diversity of species on Earth.

In the past decade, an ever-growing body of readily accessible knowledge, coupled with new tools in molecular genetics and bioinformatics, have resulted in species being described with greater accuracy, in greater detail, and with information other than morphological differences. As a result of this increase in quality and content, desirable as it is, the actual process of species description has become slower, while the rate at which species are being lost to extinction has become faster. For many groups of animals, species delimitation can be challenging using solely morphological characteristics [9], [10], and can be improved by incorporating molecular data [11], [12]. This is relevant for the conservation of threatened animals because laws or programs can be implemented more effectively when the existence of a species or population is formally described.

DNA barcoding, which is a diagnostic technique that utilizes short conserved DNA sequences, has become a popular tool for a variety of studies including species identification and molecular phylogenetic inference [13], [14], [15]. Ongoing initiatives, such as ‘Barcode of Life’ (www.barcodeoflife.org), seek to identify species and create large-scale reference databases via diagnostic DNA sequences using a standardized approach to accelerate taxonomic progress. While projects utilizing standard molecular markers have grown in popularity in the last decade, a fundamental challenge remains in transporting biological material to a site that can carry out the DNA sequencing. For instance, field biologists are often tasked with collecting samples in remote areas and subsequently transporting material to a geographically distant facility for long-term storage and molecular analyses. Furthermore, complex and overwhelming regulations can impede biological research in biodiverse countries, and can make it challenging to export material out of the country of origin [16], [17]. Additionally, many research institutions in developing parts of the world do not have access to conventional sequencing technologies within the country, further limiting identification options. This is the case for Ecuador, where most laboratories ship their samples internationally to be sequenced, often creating a delay of weeks to months between tissue collection and the availability of the sequence data. Performing genetic analyses on site or at a nearby facility within the country can help to avoid project delays and decrease the risk of sample quality decline associated with extensive transport. Now it has become possible to take portable lab equipment to remote regions, perform *in situ* experiments, and obtain genetic information relevant for biological studies and conservation policies in real-time.

The MinION (Oxford Nanopore Technologies) is a recently developed nanopore-based DNA sequencing platform. This technology has several advantages over traditional sequencing technologies, including long-read output, low initial startup costs relative to other commercial sequencers, portability, and rapid real-time analysis (reviewed by [18], [19]). Due to its small size (10 x 3.2 x 2 cm), light weight (90 grams) and ease of power and data transfer (a single USB connection to a standard laptop computer), the MinION has emerged as a valuable tool for portable sequencing projects. This device has been applied in remote sites outside of conventional labs including West Africa to monitor the 2014-2015 Ebola outbreak [20] and Brazil for Zika virus outbreak surveillance [21], [22]. It has also been applied in the Arctic to sequence microbial communities [23], [24], on military training grounds in Alberta, Canada to diagnose pathogenic bacteria [25] and in Snowdonia National Park for shotgun genomic sequencing of closely-related plant species [26]. The MinION has even been run aboard the International Space Station to evaluate performance off-Earth [27], however, the sequencing runs were performed using DNA libraries pre-prepared in a standard laboratory environment, whereas preparing samples outside of a lab with limited infrastructure presents additional challenges. Indeed, nanopore sequencing appears to hold promise for a variety of molecular experiments in the field.

Scientists have mused over the possibility of a portable method for DNA barcoding for over a decade [28], [15] and in this study our goal was to determine if the steps involved in barcoding, including real-time sequencing with the MinION, could be carried out entirely during a field expedition. Here we report conducting molecular experiments in a remote rainforest, including DNA extraction, PCR amplification of DNA barcodes, and real-time nanopore sequencing of fauna for rapid species identification. We specifically targeted DNA barcodes with existing reference databases because they are the standard approach in molecular biodiversity studies, and allowed us to rapidly produce genetic data for the identification of several animal taxa by multiplexing. Our field site was situated in a remote tropical rainforest and did not offer the commodities of a sophisticated laboratory environment, including consistent power sources or internet access. Furthermore, we restricted our laboratory equipment to reasonably affordable technologies. We did this for two reasons, (a) researchers in the field of molecular ecology may have limited funds for biodiversity research projects and (b) to test technologies that are affordable for research facilities in developing countries. We assessed the feasibility for *in situ* genetic sequencing of reptiles and amphibians for rapid species identification, using a portable laboratory fitting within a single backpack, at one of the world’s most imperiled biodiversity hotspots, the Ecuadorian Chocó rainforest (Fig. 1). We demonstrate that portable DNA amplicon sequencing with the MinION allows rapid, accurate, and efficient determination at the species level under remote tropical environmental conditions, as well as quick turnaround time for DNA barcodes of undescribed and threatened species at a research facility within the country.

**Figure 1.**
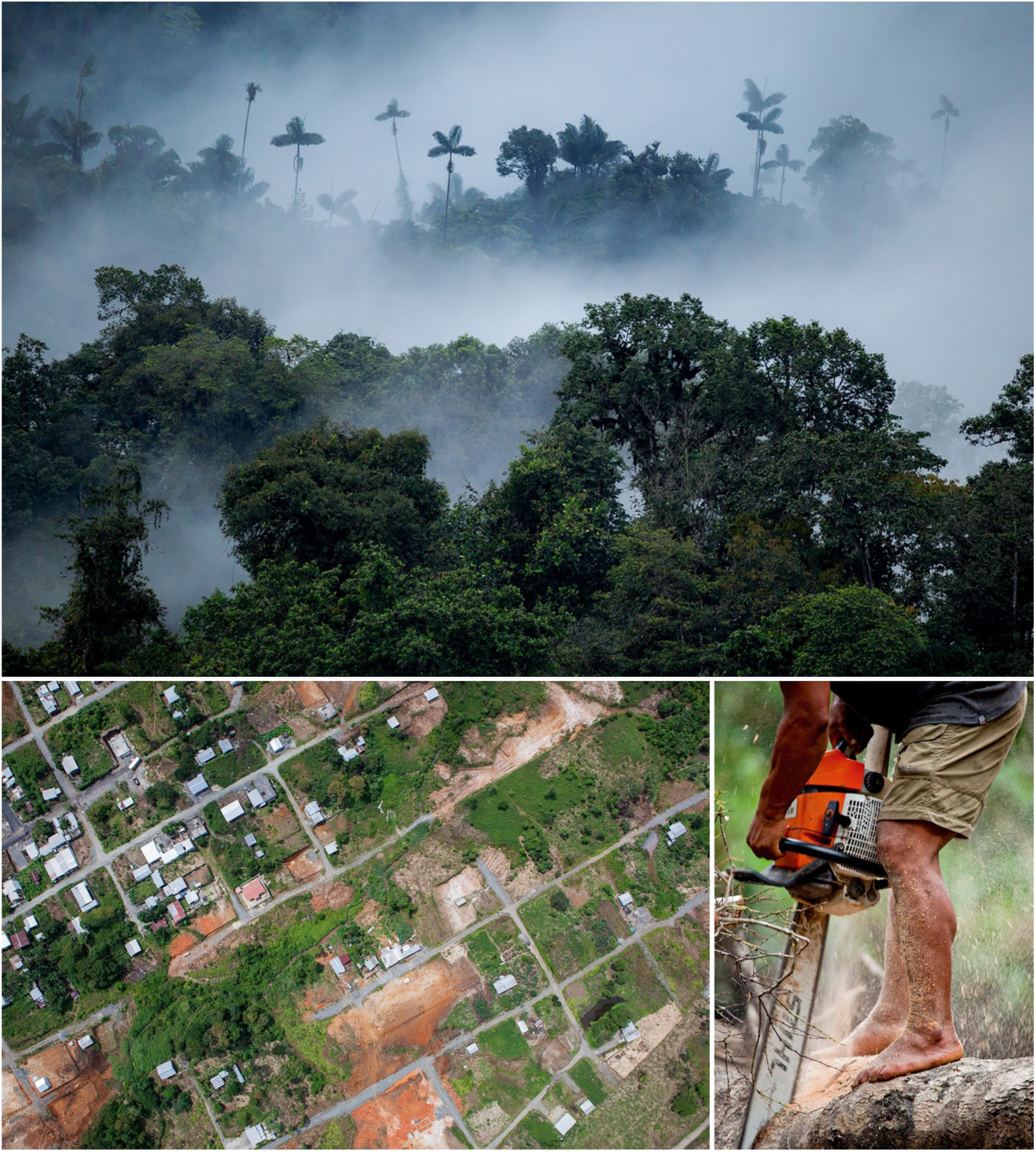
Site where field-based nanopore research was conducted within the Chocó biogeographical region in Ecuador, which is one of the world’s 25 biodiversity hotspots. This area has experienced one of the highest rates of deforestation in the country and is considered a global conservation priority.

## Methods

### Site, sampling, digital photos, tissue collection

We performed all field-based research in the Canandé Reserve (Fig. 1, 0.52993 N, 79.03541 W, 594 m), a 2000 ha. protected area, owned by Jocotoco Foundation (http://www.fjocotoco.org/canandeacute1.html) in Esmeraldas province in Northwestern Ecuador. The reserve is located in the Chocó ecoregion and is approximately 6 hours by car, depending on road conditions, from the city of Quito. The majority of organisms sampled in this study were located by space-constrained visual examination of ground-level substrates [29]. The remaining individuals were detected by turning over logs, rocks, and other surface objects. All specimens included in the genetic analyses were morphologically identified based on [30] and [31]. The sample (a tadpole, CJ 7191) of *Atelopus ignescens* was provided by the Museum of Centro Jambatu, Ecuador and was preserved in ethanol 95%. We took vouchers for all samples collected and processed in the field. These were deposited at the Museo de Zoología of the Universidad Tecnológica Indoamérica (MZUTI 5375 *Bothriechis schlegelii*, MZUTI 5383 *Lepidoblepharis* aff. *grandis*. (Gecko 1), MZUTI 5384 *Lepidoblepharis* aff. *buchwaldi*. (Gecko 2)).

### Portable laboratory equipment and set-up

The main items for portable laboratory equipment included the following: two MinION devices, a USB 3.0 cable, three SpotON flow cells (R9.5, Oxford Nanopore Technologies (ONT)), one miniPCR thermocycler (miniPCR), and a benchtop centrifuge (USA Scientific), as well as standard laboratory pipettes and sample racks (Fig. 2, Supplementary Figure 1). The MinKNOW offline software (ONT) required for operation of the MinION was installed and ran on a Windows Vaio Sony laptop with an external SSD drive (VisionTek, 240GB). All heat block and temperature cycling steps were performed using the miniPCR machine, which is a portable thermo-cycler weighing 0.45 kg. The miniPCR was programmed via an application on the laptop and powered by an external battery (PowerAdd). The total amount of equipment could fit in one carry-on backpack; a full list of laboratory hardware is provided as Supplementary Table 1. Reagents for sequencing required frozen transport from the US, which was attained by use of packaging with cold packs in a Styrofoam box and was later transferred to a plastic cool box with further cold packs upon arrival to Quito, Ecuador. MinION flow cells require storage at +2-8°C and were therefore transferred in a food storage container with chilled cold packs. At the field site, reagents and supplies were stored inside a local refrigerator and freezer.

**Figure 2.**
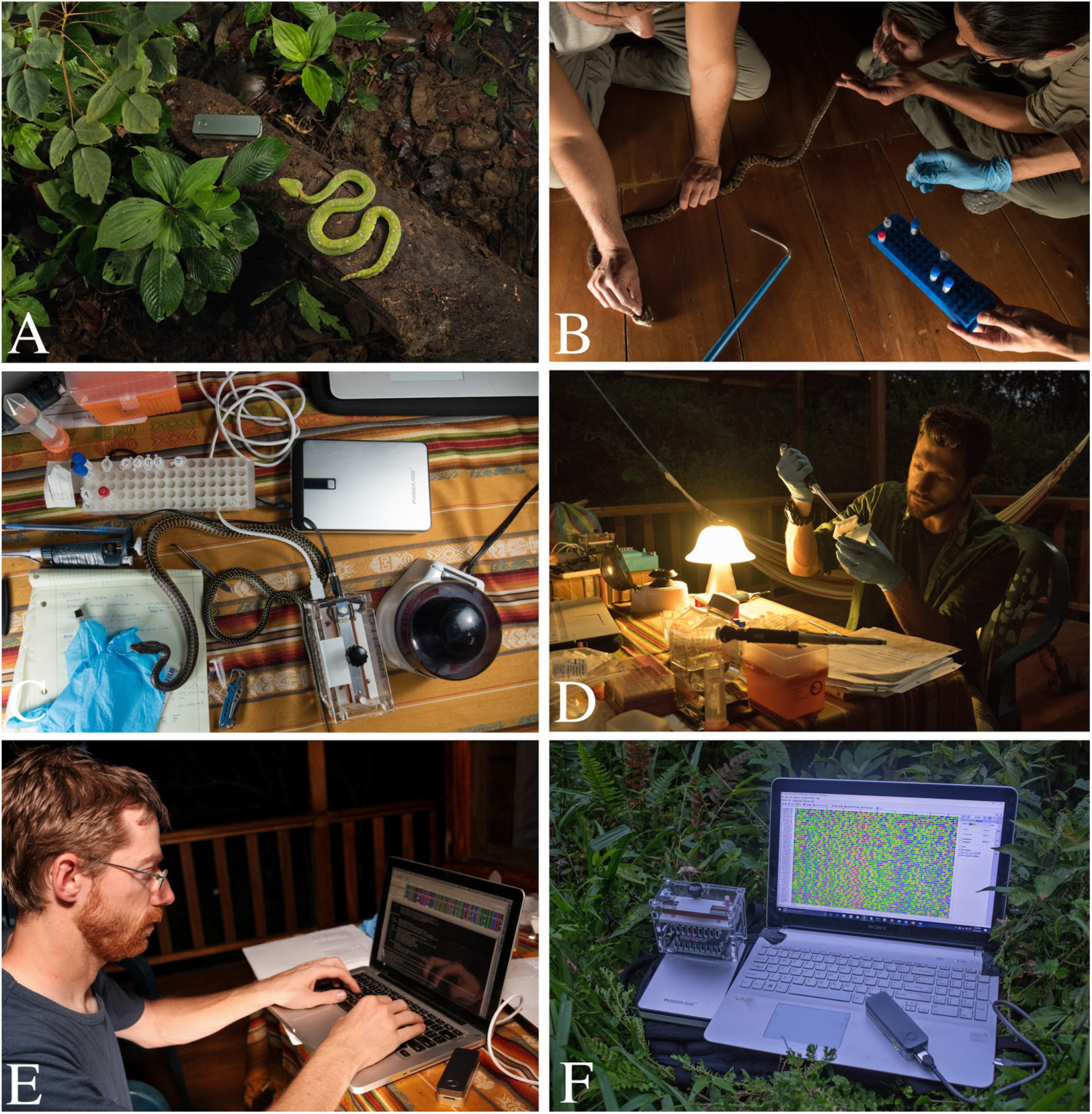
Process of nanopore sequencing in the Ecuadorian Chocó rainforest. A) Sampling endemic fauna; eyelash viper next to MinION. B) Extraction of blood or tissue samples. C) DNA extraction using the DNeasy kit and benchtop centrifuge, and PCR amplification with the MiniPCR. D) Oxford nanopore library preparation of DNA barcodes. E) Bioinformatic processing of nanopore data in the field. F) Primary equipment used in portable sequencing, left to right: MiniPCR sitting atop Poweradd external battery, MinION plugged into a Windows laptop displaying Geneious Pro software of raw nanopore data.

### Molecular techniques

Genomic DNA was extracted from fresh blood or tissue samples stored in 95% ethanol using either the DNeasy Blood & Tissue Kit (Qiagen, Hilden, Germany) according to manufacturer’s protocol and eluted in 100 μl ddH2O or a modified salt precipitation method based on the Puregene DNA purification kit (Gentra Systems) that involved cellular lysis with SDS and proteinase K, protein precipitation using guanidine isothiocyanate, and DNA precipitation by isopropanol. Tools for manipulating and lysing tissues were sterilized with a flame in between processing samples. We amplified the following mitochondrial DNA fragments: 16S gene using primers 16Sar-L and 16Sbr-H-R from [32], CytB gene using primers L14910 and H16064 developed by [33], and the gene coding for subunit 4 of the NADH dehydrogenase with primers ND4 developed by [34]. All PCR primers contained universal tailed sequences for the Oxford Nanopore Technologies barcoding kit (Supplementary Table 2). We used the ONT PCR Barcoding Kit that allows up to 12 different libraries (barcodes 1-12) to be combined and loaded onto a single flow cell at the same time. PCR reactions contained approximately 1 μl of PCR product, 2.5 μl 10X PCR buffer, 1 μl 25mM MgCl_2_, 200 μM dNTP mix, 0.2 μM of each primer and 0.25 Platinum Taq DNA Polymerase (Thermo Fisher Scientific) in a 25 μL total volume. All samples for the first PCR run were amplified on the same miniPCR under the following settings: initial denaturation: 94**°**C for 2 minutes, 35 cycles of denaturation at 94**°**C for 45 seconds, annealing at 56**°**C for 60 seconds extension for 72**°**C for 60 seconds, and a final extension of 72**°**C for 120 seconds. Then a second round of PCR was carried out, including 2 μl of ONT PCR Barcode, 2 μL of first-round PCR product, 41 μl H20, and 50 μl PCR reaction mix (0.5 μl Taq DNA polymerase, 1 μL dNTP mix, 2 μL MgCl2, 41 μL H2O). The second round of PCR barcode conditions were modified based on ONT protocol for the Platinum Taq polymerase used in this study as follows: initial denaturation at 95**°**C for 3 minutes, 15 cycles of denaturation at 95**°**C for 15 seconds, annealing at 62**°**C for 15 seconds, extension at 72**°**C for 60 seconds, and final extension at 72**°**C for 120 seconds. For verification of samples sequenced in the field, PCR products were subsequently cleaned with Exonuclase I and Alkaline Phosphatase (Illustra ExoProStar by GE Healthcare) at the Universidad Tecnológica Indoamérica (UTI) in Quito and sent to Macrogen Inc (Korea) for Sanger sequencing. All PCR products were sequenced on an ABI3730XL sequencer in both forward and reverse directions with the same primers that were used for amplification. The created sequences were deposited in GenBank (and will be available upon publication). All original Sanger and MinION generated consensus sequences can be found in Additional File 1.

### MinION sequencing

DNA library preparation was carried out according to the 1D PCR barcoding amplicons SQK-LSK108 protocol (Oxford Nanopore Technologies). Barcode DNA products were pooled with 5 μl of DNA CS control and an end-repair was performed (NEB-Next Ultra II End-prep reaction buffer and enzyme mix, New England Biolabs), then purified using AMPure XP beads. Adapter ligation and tethering was then carried out with 20 μl Adapter Mix (ONT) and 50 μl of NEB Blunt/TA ligation Master Mix (New England Biolabs). The adapter ligated DNA library was then purified with AMPure beads, followed by the addition of Adapter Bead binding buffer (ONT), and finally eluted in 15 μl of Elution Buffer (ONT). Each R9 flow cell was primed with 1000 μl of a mixture of Fuel Mix (Oxford Nanopore Technologies) and nuclease-free water. Twelve microliters of the amplicon library was diluted in 75 μL of running buffer with 35 μL RBF, 25.5 uL LLB, and 2.5 μL nuclease-free water and then added to the flow cell via the SpotON sample port. The “NC_48Hr_sequencing_FLO-MIN107_SQK-LSK108_plus_Basecaller.py” protocol was initiated using the MinION control software, MinKNOW (offline version provided by ONT).

### Bioinformatics

The commands used can be found in the Supplementary Materials and Methods section. To retrieve the nucleotide sequences from raw signal data generated by the MinKNOW software, we used Albacore 1.2.5 (https://github.com/dvera/albacore) for base calling and de-multiplexing of the ONT barcodes. The FAST5 files were then converted to fastq files using Nanopolish [35]; https://github.com/jts/nanopolish). We then filtered the raw reads for quality (score of >13) and read length (> 200bp) using Nanofilt (https://github.com/wdecoster/nanofilt), and generated consensus sequences using both reference-based mapping and *de novo* assembly. For the reference-based mapping we used BWA 0.7.15 [36]; https://github.com/lh3/bwa/releases) to align the reads to the reference, samtools 1.3 [37] to process the mapping file, and ANGSD [38], to call the consensus sequence. The *de novo* assembly of each amplicon was carried out using Canu [39], https://canu.readthedocs.io), with parameters fitting for our application. Given that we used short amplicons for the assembly we set the minimum read length to 200bp and the minimum overlap to 50bp. We subsequently extracted the consensus sequences using tgStoreDump. After the consensus calling (for both methods) we mapped the reads back to the consensus sequence (using BWA mem and samtools as described above) and polished the sequencing using Nanopolish [35]. Adapters were removed using Cutadapt [40]. The consensi were then aligned to the Sanger sequences of the same amplicons to investigate the quality of the consensus sequences generated from MinION reads using SeaView [41] and AliView [42]. Sanger sequencing reads were edited and assembled using Geneious R10 software [43] and mapping files inspected by eye using Tablet [44].

We further tested the impact of coverage on the consensus accuracy by randomly subsampling three sets of 30, 100, 300 and 1,000 reads, respectively for the eyelash palm pitviper and gecko 1. Subsampling was performed with famas (https://github.com/andreas-wilm/famas). These sets were assembled *de novo* and processed using the same approach we used for the full data sets (see above).

We then created species alignments for all barcodes (using sequences obtained from Genbank; accession numbers can be found in the phylogenetic tree reconstructions in the Supplementary material). We inferred the best substitution model using jModelTest [45] and reconstructed their phylogenetic trees using the maximum likelihood approach implemented in Mega 5 [46] with 1,000 bootstrap replicates (for bioinformatics workflow see Fig. 3). The output tree files including the Genbank Accession Numbers are provided in the supplementary material.

**Figure 3.**
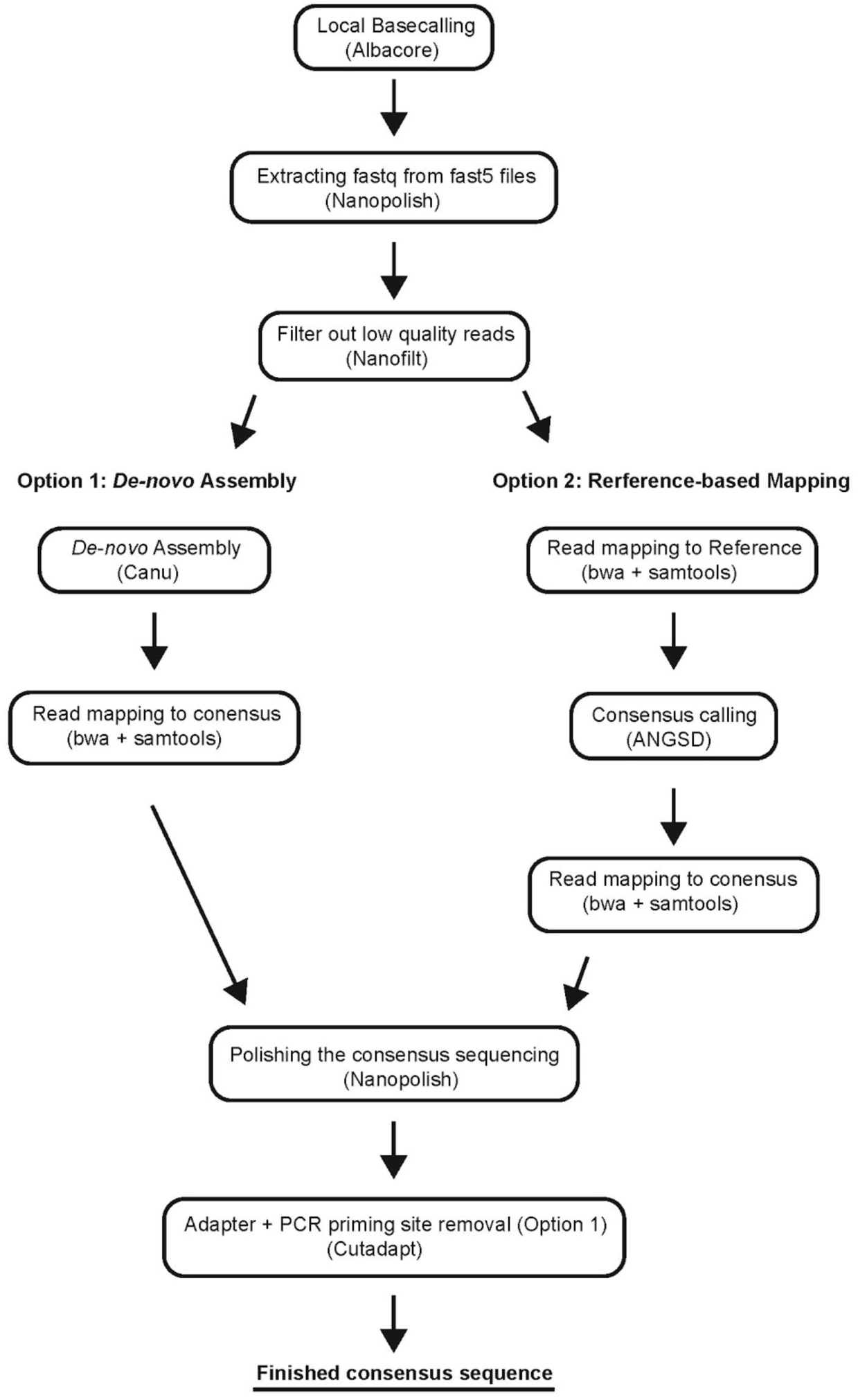
Bioinformatics workflow summarizing the steps performed during nanopore sequencing analysis with either a *de-novo* approach (left) or reference-based mapping approach (right), in order to generate a consensus sequences.

## Results

On July 11, 2017, we arrived at the field site at approximately 1500 hours and collected reptile and amphibian samples from 2000 to 2300 hours. Next, back at the field station, we extracted DNA and performed PCR amplification for 16S, CytB, and ND4 genes. On July 12, the PCR barcodes were pooled, the library was prepared, and then sequencing was initiated at approximately 1600 hours on a flow cell using the offline MinKNOW software, generating 16,663 reads after approximately two hours. The MinKNOW software was then paused in order to assess the reads generated. Within 24 hours of collecting reptiles and amphibians in the Ecuadorian Chocó, we successfully generated consensus sequences for 16S and ND4 genes of an eyelash palm pitviper (*Bothriechis schlegelii*) and 16S for the dwarf gecko (*Lepidoblepharis* sp.; gecko 1). The CytB gene was not successfully sequenced, which was later confirmed at UTI’s lab by lack of PCR product on a gel (Supplementary Figure 2). The field-generated sequence data was analyzed that evening on a laptop using a number of open source and custom-developed bioinformatic workflows (see Materials and Methods). Phylogenetic trees generated using the nanopore sequences and previously generated reference database yielded accurate species identification (Fig. 4 and Fig. 5).

**Figure 4.**
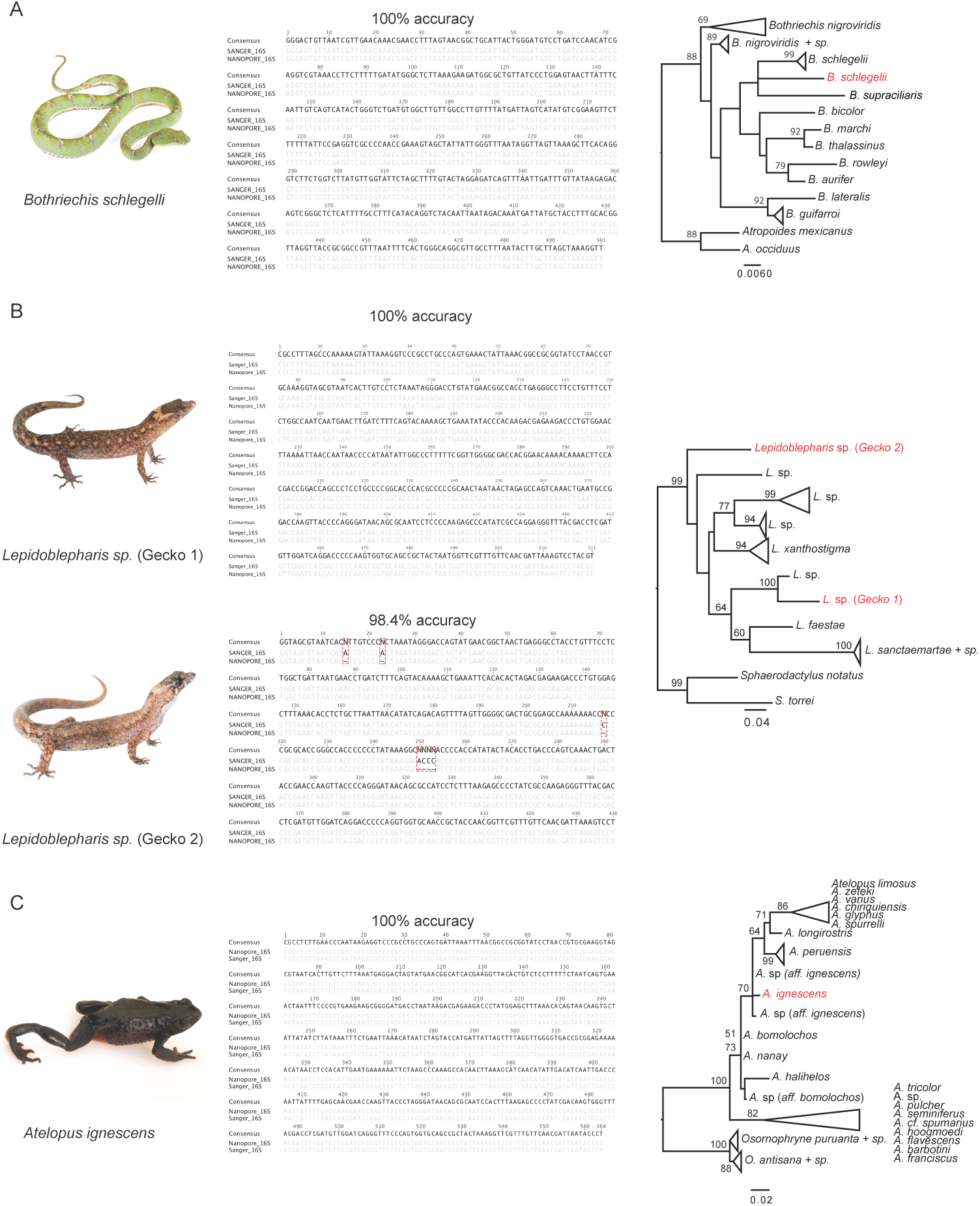
Species investigated, nucleotide alignments of nanopore and Sanger sequences comparing consensus accuracy, and Maximum Likelihood trees of 16S sequences for: A) Eyelash pitviper, Bothriechis schlegelii, B) two species of dwarf gecko, Lepidoblepharis sp, and C) the Jambato toad, Atelopus ignescens. Red labels in the phylogenetic trees indicate the sequences generated by the MinION.

**Figure 5.**
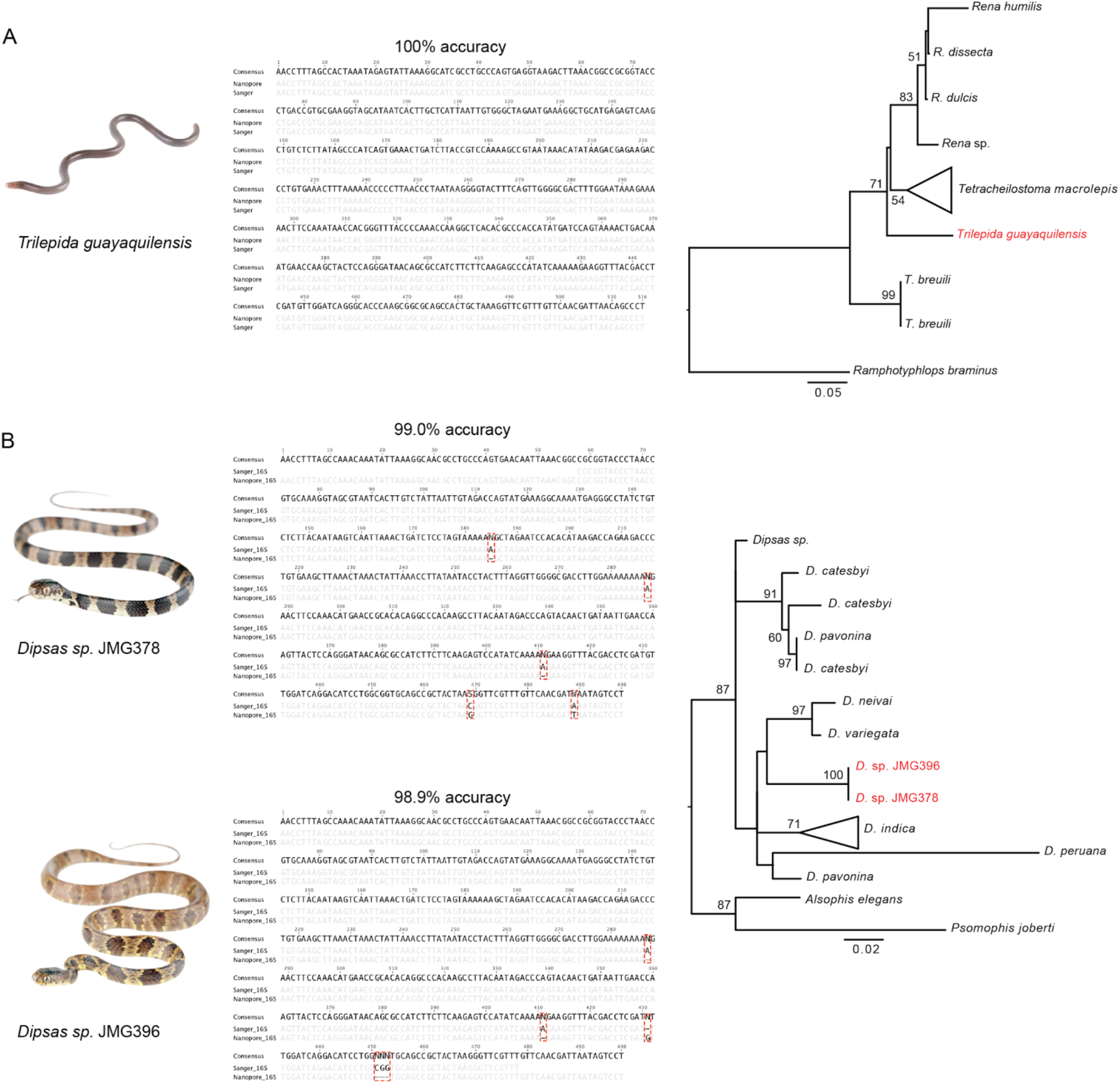
Species investigated, nucleotide alignments of nanopore and Sanger sequences comparing consensus accuracy, and Maximum Likelihood trees of 16S sequences for: A) Guayaquil blind snake, *Trilepida guayaquilensis* and B) two species *Dipsas* snakes. Red labels in the phylogenetic trees indicate the sequences generated by the MinION.

Upon returning to UTI’s lab in Quito, we created one additional DNA barcode library with new samples. With our remaining flow cell, we were interested in quickly generating genetic information for (a) additional specimens that were collected during our field expedition (gecko 2), (b) undescribed species collected the week before our expedition (Genera: *Dipsas* and *Sibon*), (c) an endangered species that would have been difficult to export out of the country (Jambato toad), (d) a rare species lacking molecular data (Guayaquil blind snake), and (e) combinations of barcoded samples through multiplexing (for the eyelash palm pitviper and gecko 1).

Initially, this second sequencing run appeared to perform well. However, after using Albacore to demultiplex the reads, we determined the adapter ligation enzyme likely degraded because the output primarily consisted of adapter sequences (Supplementary Figure 3). Nevertheless, we were able to generate consensus sequences for 16S of the Jambato toad, the two *Dipsas* species, the dwarf gecko, and the Guayaquil blind snake (Fig. 4 and Fig. 5).

The pore count of the flow cells appeared to be unaffected by travel conditions, as indicated by the multiplexer (MUX) scan, an ONT program that performs a quality check by assessing flow cell active pore count. The first run in the field had an initial MUX scan of 478, 357, 177, and 31, for a total of 1,043 active pores and after approximately two hours of sequencing the flow cell generated 16,484 reads. The second flow cell ran at UTI had a MUX scan of 508, 448, 277, and 84, for total of 1,317 active pores and the run produced 21,636 reads within two hours. This is notable since this run was performed 8 days after arriving in Ecuador and the flow cell was stored at suboptimal conditions on site and during travel.

We were unable to confidently determine if PCR bands were present in the field by running a gel with SYBR Safe DNA Gel Stain (Thermo Fisher Scientific) and a handheld ultraviolet flashlight. Therefore, the presence or absence of PCR product and size was later determined by gel electrophoresis and quantified by a Quantus Fluorometer (Promega) at UTI. Amplification for 16S and ND4 was successful for all samples, but amplification of CytB was unsuccessful, perhaps due to suboptimal PCR settings, as samples were run concurrently due to the limitation and time-constraint of having only one miniPCR machine available (Supplementary Figure 2). While the ONT protocol calls for equimolar ratios of pooled PCR product, we did not have an accurate way of quantifying DNA in the field and as such had an overrepresentation of 16S sequences, likely due to PCR bias. On future field expeditions, an inexpensive device such as the bluegel DNA electrophoresis (produced by miniPCR) can be used to assess DNA and PCR products.

### Sequencing and Bioinformatics

#### Eyelash Palm Pitviper (Bothriechis schlegelii)

The eyelash palm pitviper (*B. schlegelii*) is an iconic venomous pitviper species found in mesic forests of Central and northwestern South America [31]. The individual was captured on the evening of the 11th of July 2017 and sequenced on the MinION the following evening. We obtained 3,696 reads for the 16S fragment, 65 reads for CytB, and 94 for ND4. The 16S reads showed an average length of 655bp including the sequencing adapters. The best contig created by Canu was based on 55 reads, to which 3,695 reads mapped for the polishing step. The consensus sequence was 501bp and showed a 100% nucleotide match to the respective Sanger sequence. For this species, we did not find any differences between the *de novo* and the reference-based mapping consensus sequences (generated by mapping against a reference from the same species). The individual clusters with all other *B. schlegelii* and *B. supraciliaris* (formerly recognized as a subspecies of *B. schlegelii*) sequences in the phylogenetic tree (Fig. 4A). While the CytB *de novo* assembly did not succeed (no two reads assembled together), the best supported contig for ND4 (864bp) was based on 50 sequences and achieved an accuracy of 99.4% after polishing (using 95 reads that mapped to the *de novo* consensus).

#### Dwarf Geckos (Genus: Lepidoblepharis)

Dwarf geckos (genus: *Lepidoblepharis*) are small bodied leaf litter geckos found in Central and South America. Dwarf geckos can be difficult to identify in the field and it is suspected that there are several cryptic species within this genus in Ecuador. We captured two individuals on the evening of the 11th of July 2017, and because the two geckos differed in the shape and size of the dorsal scales (Fig. 3B) and were difficult to confidently identify by morphological characters, we decided to investigate them further with DNA barcoding.

#### Gecko 1 (*Lepidoblepharis aff. grandis)*

Gecko 1 was included in the first sequencing run in the field. We obtained 4,834 reads for the 16S fragment, 63 reads for CytB, and 76 for ND4. The consensus sequence (522bp) for this individual showed a 100% nucleotide match to the respective Sanger sequence. We then performed reference-based mapping using *L. xanthostigma* (Genbank accession: KP845170) as a reference and the resulting consensus had 99.4% accuracy. We found three insertions compared to the Sanger and the *de novo* consensus sequences (position 302: G and 350-351: AA). Next we attempted assemblies for CytB and ND4. While the assembly for the CytB reads failed, we were able to assemble the ND4 reads. However, the polished consensus sequence showed a relatively high error rate compared to the Sanger sequence (92.1% accuracy). We then blasted all ND4 reads against NCBI. For ND4 we found 8 sequences to blast to ND4 from squamates, 4 to 16S (3 to a viper and 1 to a gecko), 3 to the positive control, 10 very short hits (negligible hits), and 46 to find no blast hit. Interestingly, while only 8 reads were hits for ND4 from squamates, 72 reads mapped to the consensus of the *de novo* assembly. The higher error rate can thus be explained by the fact that contaminant reads were used to assemble and correct consensus. The *de novo* assembled consensus showed an accuracy of 91.7% compared to 92.1% for the polished sequence.

#### Gecko 2 (*Lepidoblepharis aff. buchwaldi*)

Gecko 2 was included in the second sequencing run at UTI. We generated 325 reads (for more information see discussion on the possible issue with the adapter ligation enzyme). After filtering for read quality and assembly, we found the best contig to be supported by 30 reads. Out of the 325 barcoded reads, we found 308 to map to the consensus. After running Nanopolish, we found it to match 98.4% to the Sanger sequence. All of the observed differences were indels (mostly 1 bp, but also one 4 bp indel; positions: 15, 23, 217 and 250-253, respectively, Fig. 4B). Positions 15 and 23 show an A in the reference, which is not found in the nanopore consensus (filtered or unfiltered, and polished or not polished). Position 217 is a C in the Sanger reference. None of the consensi for the nanopore data showed the C. This error can potentially be explained as it lies within a 6 bp cytosine homopolymer (see Lu et al., 2016). Interestingly, we saw only a 1bp mismatch instead of the 4bp indel at position 250-253 in the filtered, but not polished nanopore consensus sequence. After polishing all sequences (filtered or unfiltered) showed the 4bp indel. We next applied reference based mapping (same protocol and reference as for gecko 1). The resulting consensus sequence showed an accuracy of 97.9%. Phylogenetic tree reconstruction shows that gecko 1 and gecko 2 are clearly two distinct species (see Fig. 4B).

#### Jambato toad (Atelopus ignescens)

Laboratory processing and sequencing for *Atelopus ignescens* was carried out in the lab at UTI using a preserved tadpole sample. We obtained 503 reads for this species. The best supported *de novo* assembled contig was based on 56 reads. We then mapped the reads back to this contig for the polishing step, which resulted in 491 mapped reads. However, while the total coverage was 434x for the segment, the average coverage was only 212x. The discrepancy can be explained by a high percentage of reads that exclusively consisted of adapter sequences (probably caused by inefficient adapter ligation; see Discussion section; Supplementary Figure 3). The resulting sequence fits 100% to the respective Sanger sequence (Fig. 4C). We next used the reference-based approach to construct a consensus sequence, using *Atelopus hoogmoedi* (Genbank accession: EU672974) as a reference and the consensus achieved an accuracy of 100% after polishing. The phylogenetic tree reconstruction clusters our sequence with samples described as *A*. sp *aff. ignescens*.

#### Guayaquil blind snake (Trilepida guayaquilensis)

The Guayaquil blind snake (*Trilepida guayaquilensis*) belongs to the family of Slender blind snakes (Leptotyphlopidae). This family is found in North and South America, Asia, and Africa. They are fossorial snakes adapted to life underground. The Guayaquil blind snake was only known from one individual described in 1970 and is suspected to be endemic to Ecuador [47]. For a second specimen collected recently, we obtained 756 sequences. However, many of those reads were adapter sequences. The Canu *de novo* assembled sequence was generated from 16 reads. We then mapped 740 reads back to this consensus. After polishing the consensus sequence matched 100% of the Sanger generated sequence (Fig. 5A; 516bp consensus length). We further investigated the accuracy of reference based mapping for this species. We used *Trilepida macrolepis* (Genbank accession: GQ469225) as a reference, which is suspected to be a close relative of *T. guayaquilensis*. However, the resulting consensus sequence had a lower accuracy (97.7%) compared to the *de novo* assembled consensus (100%). Our sequence is sister to the clade comprising *Trilepida macrolepis* and all *Rena* species in the phylogenetic tree.

#### Dipsas snakes (Genus: Dipsas)

*Dipsas* are non-venomous New World colubrid snakes that are found in Central and South America (Cadle 2005). Here we included two specimens collected one week prior to our expedition, which are suspected to be undescribed species.

#### *Dipsas sp*. (JMG378)

We generated 779 reads for *Dipsas* (JMG378). The best supported contig of the Canu *de novo* assembly (498bp consensus length) was based on 59 reads and matched the corresponding Sanger sequence to 99% after polishing (Fig. 5B). Three out of 5 mismatches were indels in poly-A stretches (position: 185, 287, 411). The remaining two mistmachtes are a C to G at position 469 and a T to A at position 489 for the nanopore compared to the Sanger sequence. Interestingly, the reference-based consensus sequence (using *Dipsas* sp., GenBank accession: KX283341 as a reference) matched the Sanger sequence to 99.4% after polishing. We generated 816 reads for the CytB barcode. However, *de novo* assembly was not successful as none of the reads actually belonged to CytB.

#### *Dipsas sp*. (JMG396)

We generated 487 reads for *Dipsas* (JMG396). Sequences with a quality score of >13 were retained resulting in 193 sequences. The best supported contig of the Canu *de novo* assembly was based on 59 reads (498bp consensus length). After polishing the consensus sequence matched the corresponding Sanger sequence to 98.9% (Fig. 5B). The first two mismatches are typical nanopore errors, namely indels in poly-A stretches (positions: 287, 411). The nanopore sequence shows an insertion of a single G compared to the Sanger sequence as position 431. The last mismatch is a three base pair deletion compared to the Sanger sequence (positions: 451-453). The reference-based consensus (using *Dipsas* sp., GenBank accession: KX283341 as a reference) achieved a 98.4% match after polishing. We generated 1,077 reads for the CytB barcode. Again, *de novo* assembly was not successful as none of the reads actually belonged to CytB. The two *Dipsas* specimens clustered together in the phylogeny. They are sister to the clade comprising *D. neivai* and *D. variegata*. However, this part of the phylogeny shows low support (bootstraps < 50).

#### Sibon snake (Genus: Sibon)

Sibon snakes are found in northern South America, Central America and Mexico [48]. We generated 339 reads for the 16S barcode of this species. However, we were not able to create a consensus sequence for this barcode, as almost all the reads were adapter sequences (all but 11 reads). Furthermore, we generated 1,425 reads for the CytB barcode but were not able to create a consensus sequence.

#### Subsampling

We further investigated the read depth needed to call accurate consensus sequences using our approach. We used the eyelash palm pitviper and gecko 1 to test subsampling schemes, since we obtained thousands of reads for these samples. We randomly subsampled to 30, 100, 300 and 1,000 reads (in three replicates; see Supplementary Table 3). For the eyelash palm pitviper we achieved accuracies ranging from 99.4% to 99.8% using only 30 reads, 99.6% to 100% using 100 reads, 99.8% for 300 reads and 99.8% to 100% for 1,000 reads. For gecko 1 we achieved even better accuracy overall, with 30 reads ranging from 99.4% to 99.8%, 100 reads from 99.8% to 100%, all 300 reads sets achieved an accuracy of 100% and for 1,000 reads all but one set (99.8%) achieved 100% accuracy.

#### Multiplexing

We further sequenced multiplexed barcodes (16S and ND4) for the eyelash palm pitviper and gecko 1. However, we did not obtain reads for this sample from sequencing run 2, most likely due to the adapter ligation issues. We thus generated artificial multiplexes for the eyelash palm pitviper pooling random sets of 1,000 16S reads with all 96 ND4 reads to investigate the performance of the *de novo* assembly using multiplexed samples. We assembled the reads *de novo* and processed them using the same approach as discussed above. In all three cases we found the first two contigs of the canu run to be 16S and ND4 contigs. After polishing the 16S consensus sequences achieved a 99.8% accuracy (all three assemblies showed a deletion in a stretch of four T’s compared to the Sanger sequence) and the ND4 sequences a 99.4% accuracy. All errors, but one (which shows a T compared to the C in the Sanger sequence), in ND4 are deletions in homopolymer stretches.

## Discussion

In this project, we investigated the feasibility of molecular species identification in a remote tropical field location. Below we discuss different aspects of the project such as performance, conservation implications, significance for local resource building, as well as outlooks and improvements for future development.

### Performance in the field

Our objective was to employ a portable laboratory in a rainforest to identify endemic species with DNA barcoding in real-time. Our protocols resulted in successful DNA extraction, PCR amplification, nanopore sequencing, and barcode assembly. We observed that the MinION sequencing platform performed well in the field after extended travel, indicating the potential for nanopore-based sequencing on future field expeditions. Although we demonstrate that the successful molecular identification of organisms in a remote tropical environment is possible, challenges with molecular work in the field remain. Our field site was provided with inconsistent electrical power, but still allowed us to use a conventional small centrifuge for several steps of DNA extraction and to power a refrigerator for storage of flow cells and some of the reagents, although temperatures were likely suboptimal. Lack of electrical supply can impede adequate storage of temperature-sensitive reagents for extended periods of time. Our experiments were performed during a relatively short field trial, with 10 days being the longest time period that reagents were kept at inconsistent freezing temperatures. It is uncertain how well nanopore kit reagents or flow cell integrity would endure over longer periods without consistent cooling temperatures, and we suspect the adapter ligation enzyme was compromised during our second nanopore run, as demultiplexing led to a majority of barcode adapters in each folder (Supplementary Figure 3). We used an external SSD drive with 240 GB space to store raw data generated by MinKNOW. Due to overheating of the external drive, we placed ice packs underneath the USB stick to maintain cooler temperatures, which appeared to maintain the run.

While the MinION sequencer fits in the palm of a hand and needs only a USB outlet to function, bioinformatic analyses can be hampered under remote field conditions, because internet access, large amounts of data storage, and long periods of time are often required for such analytical tasks. In our study, utilizing short DNA fragments with a relatively small number of samples for barcoding allowed us to perform all bioinformatic analyses in the field, but larger data outputs may require additional storage and more computational resources.

### Implications for conservation

Tropical rainforests, such as the Ecuadorian Chocó, are often rich in biodiversity, as well as species of conservation concern. The Chocó biogeographical region is one of the world’s 25 biodiversity hotspots [49] and several studies have identified the Chocó region of western Colombia and Ecuador as a global conservation priority [49], [50], [51]. We therefore chose this region for proof of principle *in situ* molecular work to highlight the importance of expediting fieldwork in order to produce genetic information of endemic fauna. Our rapidly obtained DNA barcodes allowed us to accurately identify organisms while in the field, and had an accuracy comparable to Sanger generated sequences. When samples are not required to be exported out of the country to carry out molecular experiments, real-time sequencing information can contribute to more efficient production of biodiversity reports that advise conservation policy, especially in areas of high conservation risk.

Of particular note in this study was the critically endangered harlequin Jambato toad, *Atelopus ignescens*. Although not a denizen of the Chocó rainforests, this Andean toad is a good example to demonstrate how nanopore sequencing can aid in the conservation of critically endangered species. *Atelopus ignescens* was previously presumed extinct (it is currently still listed as “extinct” on IUCN; [52] and was only recently rediscovered [53]. *Atelopus* is a species-rich genus of neotropical toads containing 96 species, most of which are possibly extinct or endangered. In Ecuador there are 11 species of *Atelopus* that are Critically Endangered (tagged as Possibly extinct; [54]). Extinctions of *Atelopus* (and other anurans) are beyond control and are increasingly exacerbated by a combination of factors including habitat loss, climate change and pathogens [55], [56], [57]. For the many endangered species that are protected by international laws and treaties, sample transport requires permits that can often be difficult to obtain, even when research is expressly aimed at conservation, resulting in project delays that can further compromise sample quality. By working within the country, under permits issued by Ministerio del Ambiente de Ecuador to local institutions, we were able to generate sequence data for the endangered harlequin Jambato toad *Atelopus ignescens* within 24 hours of receiving the tissue, whereas obtaining permits to ship samples internationally in the same time frame would have not been possible. The last confirmed record of *Atelopus ignescens* dates back to 1988, and this species was presumed to be extinct before one population was rediscovered in 2016, 28 years later. Rapidly identifying the phylogenetic affinity of populations of *Atelopus* toads could speed up conservation efforts for these animals. Namely, a better understanding of the systematics of the group facilitated by real-time sequencing could help establish species limits, geographic distributions, in-situ conservation actions and ex-situ breeding programs.

### Species identifications

It is important to note that we do not intend for rapidly-obtained portable sequence information to substitute for standard species description processes. Instead, we aim to demonstrate that obtaining real-time genetic information can have beneficial applications for biologists in the field, such as raising the interesting possibility of promptly identifying new candidate species, information which can be used to adjust fieldwork strategies or sampling efforts. As we have shown, the latter could be especially important with organisms and habitats facing pressing threat. Rapidly obtaining genetic sequence information in the field can also be useful for a range of other applications, including identifying cryptic species, hybrid zones, immature stages, and species-complexes.

Furthermore, we acknowledge that in most cases multiple loci are needed to reliably infer species position in a phylogenetic tree. DNA barcoding has been shown to hold promise for identification purposes in taxonomically well-sampled clades, but may have limitations or pitfalls in delineating closely related species or in taxonomically understudied groups [58], [59]. However, our aim in this study was to demonstrate that portable sequencing can be used in the field and that the final sequences have an accuracy needed to achieve reliable identification of a specimen. While a recent study has demonstrated a field-based shotgun genome approach with the MinION to identify closely related plant species [26], DNA barcoding already offers a robust reference database for many taxa thanks in part to global barcoding initiatives (the current Barcode of Life Data System contains 4,013,927 specimens and 398,087 Barcode Index Numbers http://ibol.org/resources/barcode-library/ as of September 2017).

Finally, while highlighting the value of real-time portable DNA barcoding in this study, we do not wish to downplay the significance of taxonomic experts, who have invaluable specialist knowledge about specific groups of organisms. Even with the advent of molecular diagnostic techniques to describe and discover species, placing organisms within a phylogenetic context based on a solid taxonomic foundation is necessary. An integrative approach utilizing molecular data and morphological taxonomy can lead to greater insight of biological and ecological questions [60]. As noted by Bik, 2017, *“There is much to gain and little to lose by deeply integrating morphological taxonomy with high-throughput sequencing and computational workflows.”*

### Bioinformatic challenges

Although nanopore sequence reads show high error rates (12-35%; [61], [18], [62]), the consensus sequences generated in this study matched the respective Sanger sequences with high accuracy, ranging from 98.4% to 100% (Fig. 4 and Fig. 5), with four out of seven sequences showing an accuracy of 100%. In order to investigate the minimum coverage needed to accurately call bases, we used different subsampling schemes. Overall, a coverage of 30 reads achieved an accuracy of 99.4 - 99.8%. With 100 reads most assemblies were 100% accurate, indicating that an excessive amount of reads is not needed to produce high quality consensus sequences. Furthermore, we applied Nanopolish to all consensus sequences. This tool has been shown to be very effective at correcting typical nanopore errors, such as homopolymer errors [35], [63]. As can be seen in section “Post-Nanopolish assembly identity” in [63], accuracy of the resulting consensus increases significantly after polishing. While, we did not measure the improvement in accuracy in our study, we did notice a high accuracy after polishing. However, as can be seen in Fig. 5B, nanopolish is not always able to accurately correct homopolymer stretches.

We further tested reference-based mapping versus *de novo* assembly, because a reference-based mapping approach may introduce bias, making it possible to miss indels. Overall, we see that consensus sequences generated using reference-based mapping have slightly lower accuracy. However, in two cases (the eyelash palm pitviper and the Jambato toad) an accuracy of 100% was achieved with reference-based mapping. Interestingly, in the case of *Dipsas* sp. (JMG378), reference-based mapping resulted in a slightly better accuracy than *de novo* (99.4% compared to 99%). In general, we recommend the use of a *de novo* assembly approach as this method can be applied even if no reference sequence is available and generally produced more accurate sequences. An alternative approach would be to generate consensus sequences by aligning the individual reads for each barcode to one another, which would not be affected by a reference bias. This method is implemented in the freely available software tool Allele Wrangler (https://github.com/transplantation-immunology/allele-wrangler/). However, at the time of submission this tool picks the first read as the pseudo reference, which can lead to errors in the consensus if this read is of low quality or an incorrect sequence. Future developments might establish this method as an alternative to *de novo* assembly algorithms, which are typically written for larger genomes (e.g. the minimum genome size in Canu is 1000bp) and can have issues with assemblies where the consensus sequence is roughly the size of the input reads (*personal communications* Adam Phillippy).

Each of our two runs showed a very high number of reads not assigned to any barcode sequence after de-multiplexing with Albacore 1.2.5 (7,780 and 14,272 for the first and second sequencing run, respectively). In order to investigate whether these reads belong to the target DNA barcodes but did not get assigned to sequencing barcodes, or if they constitute other sequences, we generated two references (one for each sequencing run) comprising all consensi found within each individual sequencing run. We then mapped all reads not assigned to barcodes back to the reference. We were able to map 2,874 and 4,997 reads to the reference for the first and the second sequencing run, respectively, which shows that a high number of reads might be usable if more efficient de-multiplexing algorithms become available. Here we used Albacore 1.2.5, an ONT software tool, to de-multiplex the sequencing barcodes. This tool in under constant development and thus might offer more efficient de-multiplexing in later versions. Alternatively, 3rd party software tools like npBarcode [64] or Porechop (https://github.com/rrwick/Porechop) can be used.

### Cost-effectiveness and local resource development

Next-generation sequencing technologies are constantly evolving, along with their associated costs. Most major next-generation sequencing platforms require considerable initial investment in the sequencers themselves, costing hundreds of thousands of dollars, which is why they are often consolidated to sequencing centers at the institutional level [65]. In this study, we used the ONT starter pack, which currently costs $1000, and includes two flow cells and a library preparation kit (6 library preparations) as well as the ONT 12 barcoding kit which is currently $250 for 6 library preparations (for a full list of equipment and additional reagents see Supplementary Table 1). Using this setup, each barcode sequence costs about $45 (a detailed cost account can be found in the Supplementary material). At this current cost, further multiplexing of samples on each flow cell is necessary to achieve a cost-effectiveness for DNA sequencing relative to other commercial options. Fortunately, ONT also offers a 96 barcode kit, currently priced at $1,700. While this is a higher upfront cost it can reduce the price for each DNA barcode sequence to about $12. This price can be further reduced when multiplexed DNA barcodes are used. The calculated prices assume a cost of $900 per flow cell (that is not part of the starter pack), which can lessen when bought in bulk ($500 per flow cell for an order of 48). On the contrary, Sanger sequencing from UTI shipped internationally for processing costs approximately $10 per sample, independent of the through-put. Thus the Oxford Nanopore MinION has the potential to be a cost-effective sequencing option for resource-limited labs, especially in developing countries without access to standard sequencing devices.

The small size and low power requirements of the MinION will likely continue to enable its evolution as a field-deployable DNA sequencing device, opening up new avenues for biological research in areas where the typical laboratory infrastructure for genetic sequencing is unavailable. With some training, in the field molecular analyses could also potentially be performed by local students or assistants, providing an opportunity for capacity building and community involvement.

### Future outlook

Technological developments in lab equipment and reagent chemistry are increasingly enabling the incorporation of genetic analyses into field projects. Several portable technologies have been used to perform molecular experiments in the field, particularly for disease diagnostics [66], [67]. Advances in lyophilized and room-temperature reagents are also promising for field applications, such as EZ PCR Master Mix [68], and loop-mediated isothermal amplification [69], [70]. A hand-powered centrifuge could also act as substitute for a standard benchtop centrifuge during DNA extraction steps. Automatic devices, such as VolTRAX (a compact microfluidic device designed to automate nanopore library preparation, ONT) and improved library construction methods may offer faster and high-throughput methods for preparing nanopore libraries in the future. As the ONT MinION evolves, it could greatly advance field researchers’ capacity to obtain genetic data from wild organisms while in the field. These technologies currently depend on reagents that require freezing, but can be used at field sites with solar or portable freezer options. Faster and more automated sample processing, as well as cost reductions, are needed for adoption in low-income settings.

Beyond short PCR-based amplicons aimed at species identification, other exciting potential applications of nanopore sequencing in the field include sequencing of entire mitochondria from gDNA samples [71] or via long-range PCR, shotgun genome sequencing [26], analysis of environmental DNA [72], [24], sequencing of direct RNA [73], [74] or cDNA to rapidly profile transcriptomes ([75], and pathogen diagnostics and monitoring (such as chytrid fungus; [76]. Rapid portable sequencing can also be applied to wildlife crime to perform species identification of animals affected by illegal trafficking, as well as serve to aid in early detection of invasive species threatening local biodiversity and agriculture, and emerging infectious diseases.

## Conclusion

Portable DNA barcoding with the MinION sequencer allows rapid, accurate, and efficient determination at the species level under remote and tropical environmental conditions. We also demonstrate that portable sequencing can allow nimble use of rapidly generating data for endangered, rare, and undescribed species at nearby facilities within the country. In the context of conservation and biodiversity science, portable nanopore sequencing can be beneficial for applications including:

i. When it is exceedingly challenging or not possible to export biological material internationally or to a facility with a conventional sequencing device. The proper permits to collect samples and carry out experiments in the location of the study are still necessary, and collaborating with local researchers is encouraged.
ii. When the material to be sequenced may be compromised during transportation conditions, or during the time in between collection and sequencing. This can be applicable to experiments involving RNA in particular, which is subject to degradation if not adequately preserved or immediately frozen.
iii. For biodiversity reports aimed at quickly generating species data to inform conservation policy decisions, especially in areas of high conservation risk.
iv. To rapidly screen and sequence pathogens, such as chytrid fungus in amphibians. Studies using the MinION in the field have been applied during epidemics, including recent outbreaks of Ebola and Zika, and can be applied to non-human pathogens as well.
v. To perform on-site identification of organisms, immature life stages, or sexes that are difficult to distinguish morphologically, such as larvae or pupae of insects, plants when they are not actively flowering such as orchids, or cryptic species.
vi. To identify organisms in the field that are difficult to locate or capture by sampling environmental DNA (eDNA).

While we live in a period of amazing technological change, biodiversity and ecosystem health are decreasing worldwide. Portable sequencing will not be a silver bullet for conservation biology, but it can be a powerful tool to more efficiently obtain information about the diversity of life on our planet. This is particularly important for many tropical rainforests under high risk of habitat loss, such as the Ecuadorian Chocó. We present, to our knowledge, the first expedition to successfully demonstrate real-time animal species identification using DNA barcode sequencing in a remote tropical rainforest. We anticipate that as portable technologies develop further, this method will broaden the utility of biological field analyses including real-time species identification, cryptic species discovery, biodiversity conservation reports, pathogen detection, and environmental studies.

## Acknowledgements

Fieldwork for this project was made possible with the support of Tropical Herping and Universidad Tecnológica Indoamérica. For granting access to the Canandé reserve, we are grateful to Martin Schaefer of Fundación Jocotoco. Funding for equipment and reagents was supported by the National Geographic Society / Waitt grant (W412-15). Laboratory work was carried out at Universidad Tecnológica Indoamérica in Quito. Research and collection permits were issued by the Ministerio del Ambiente de Ecuador (MAE-DNB-CM-2015-0017). The Jambato toad tissue was provided by the Museum of Centro Jambatu under the Ministerio del Ambiente de Ecuador project “Conservation of Ecuadorian amphibian diversity and sustainable use of its genetic resources.” We thank Oxford Nanopore Technologies for providing technical support, making the offline MinKNOW software available, and for providing two free flow cells. We also thank Hitomi Asahara and the UC Berkeley DNA Sequencing Facility for lending the benchtop centrifuge used in this study; Jared Simpson, Sergey Koren and Adam Phillippy for very helpful advice and discussion on the bioinformatic pipeline, and Ellie E. Armstrong for valuable input on the manuscript.

## Supplementary Information

**Supplementary Figure 1.**
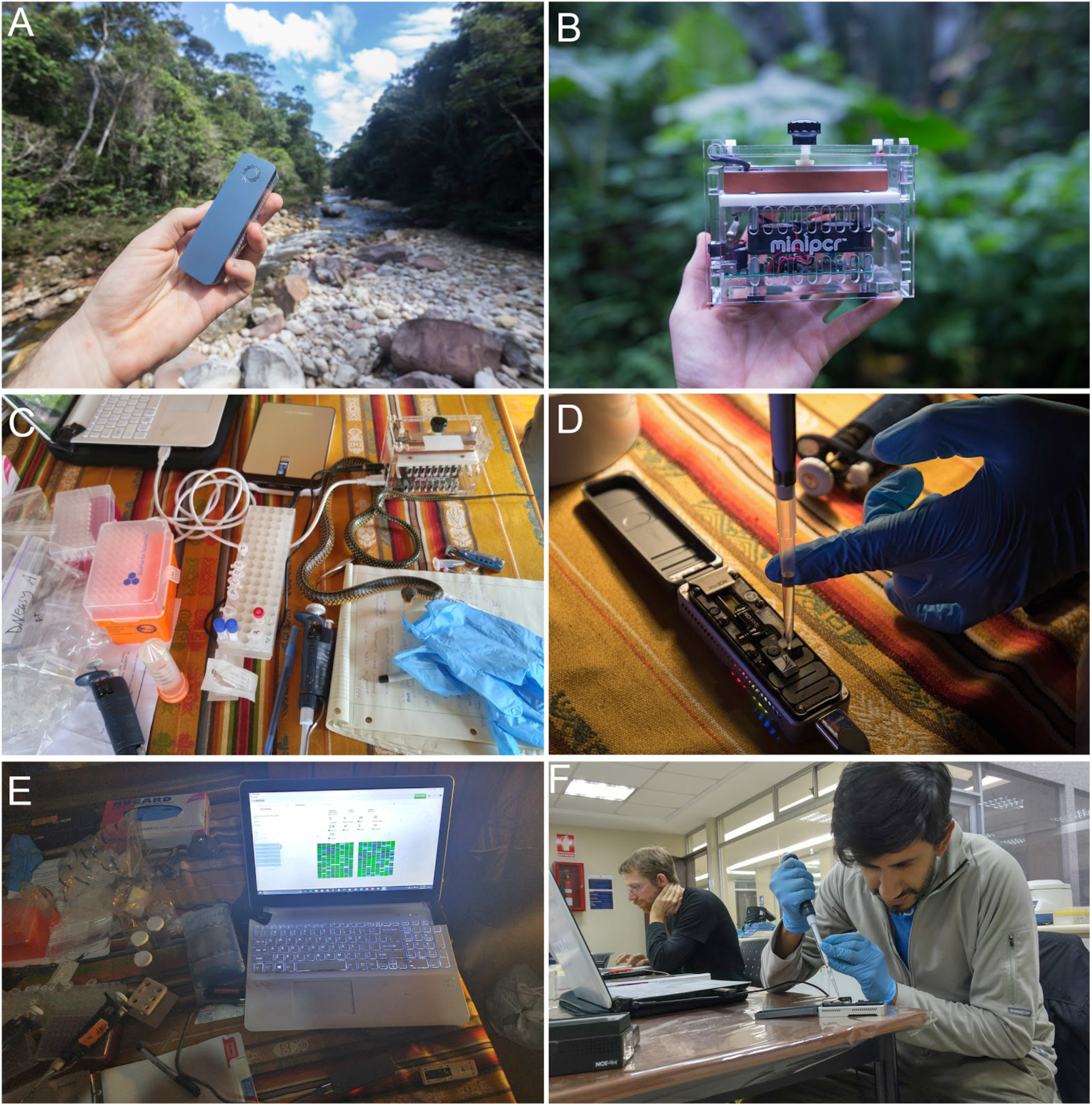
Additional images highlighting the portable lab equipment and setup used for nanopore sequencing in Ecuador. A) The handheld MinION DNA sequencer (Oxford Nanopore Technologies). B) miniPCR Thermocycler (miniPCR). C) Mobile setup for DNA extraction and PCR amplification. D) Loading the ONT flow cell in the field. E) Running the MinION using offline MinKNOW software. F) Local collaborator loading the MinION at a nearby research facility, highlighting the opportunity for capacity building and community involvement.

**Supplementary Figure 2.**
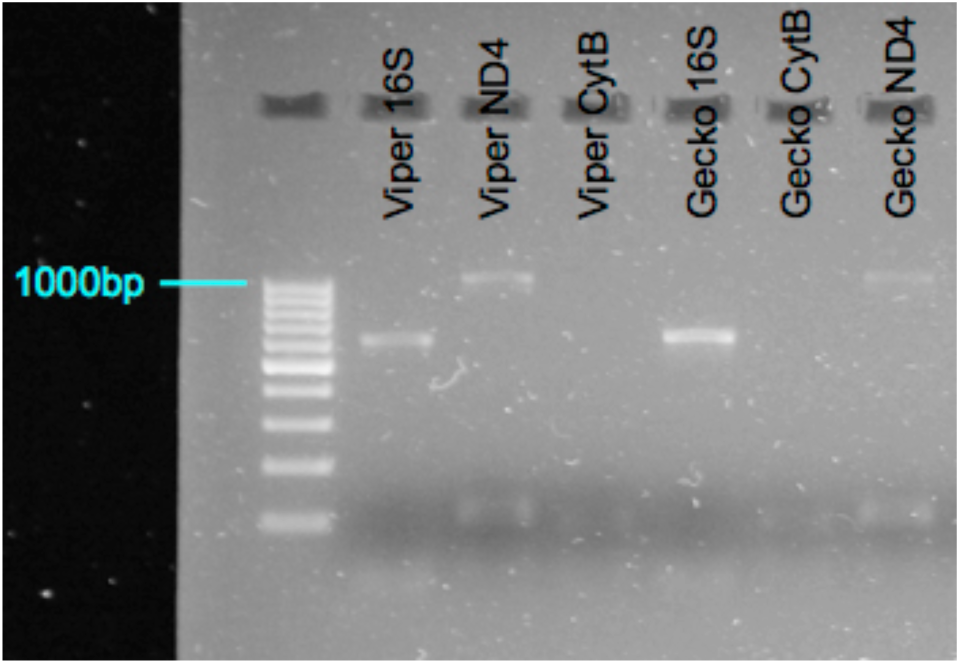
Gel of PCR product that was produced in the field using the miniPCR and imaged at UTI in Quito. Note that 16S and ND4 from samples amplified but CytB did not.

**Supplementary Figure 3.**
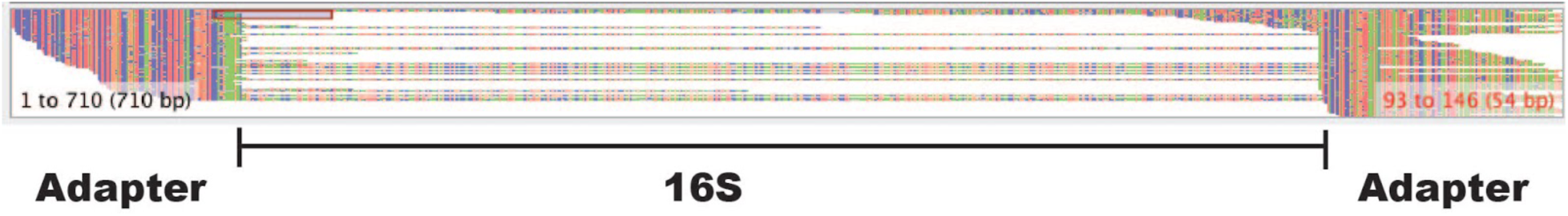
Large portion of adapter sequences contained in demultiplexed barcodes, indicating possible adapter ligation degradation for the second nanopore run at UTI.

**Supplementary Table 1.** List of equipment, consumables, and reagents used for portable nanopore sequencing in Ecuador.

| **Equipment** | **Supplier** |
| --- | --- |
| Laptop | Windows, MacBook Pro |
| MinION | Oxford Nanopore Technologies |
| Flow cell, R9.5 | Oxford Nanopore Technologies |
| miniPCR thermal cycler | miniPCR |
| External SSD Flash Drive | VisionTek |
| Benchtop centrifuge | USA Scientific |
| Pipettes, Gilson (P20, P200, P1000) | ThermoFischer |
| Magnetic rack | Promega |
| External battery source | PowerAdd |
| Gel electrophoresis chamber (MSMINI) | Biocomdirect |
| 300V electrophoresis power source | VWR International |
| Tube racks for 1.5 mL tubes | - |
| **Consumables** |  |
| Gloves | - |
| Pipette tips (P20, P200, P1000) | ThermoFischer |
| 1.5 mL Eppendorf tubes | ThermoFischer |
| 0.2 mL PCR tubes | ThermoFischer |

**Supplementary Table 2.**
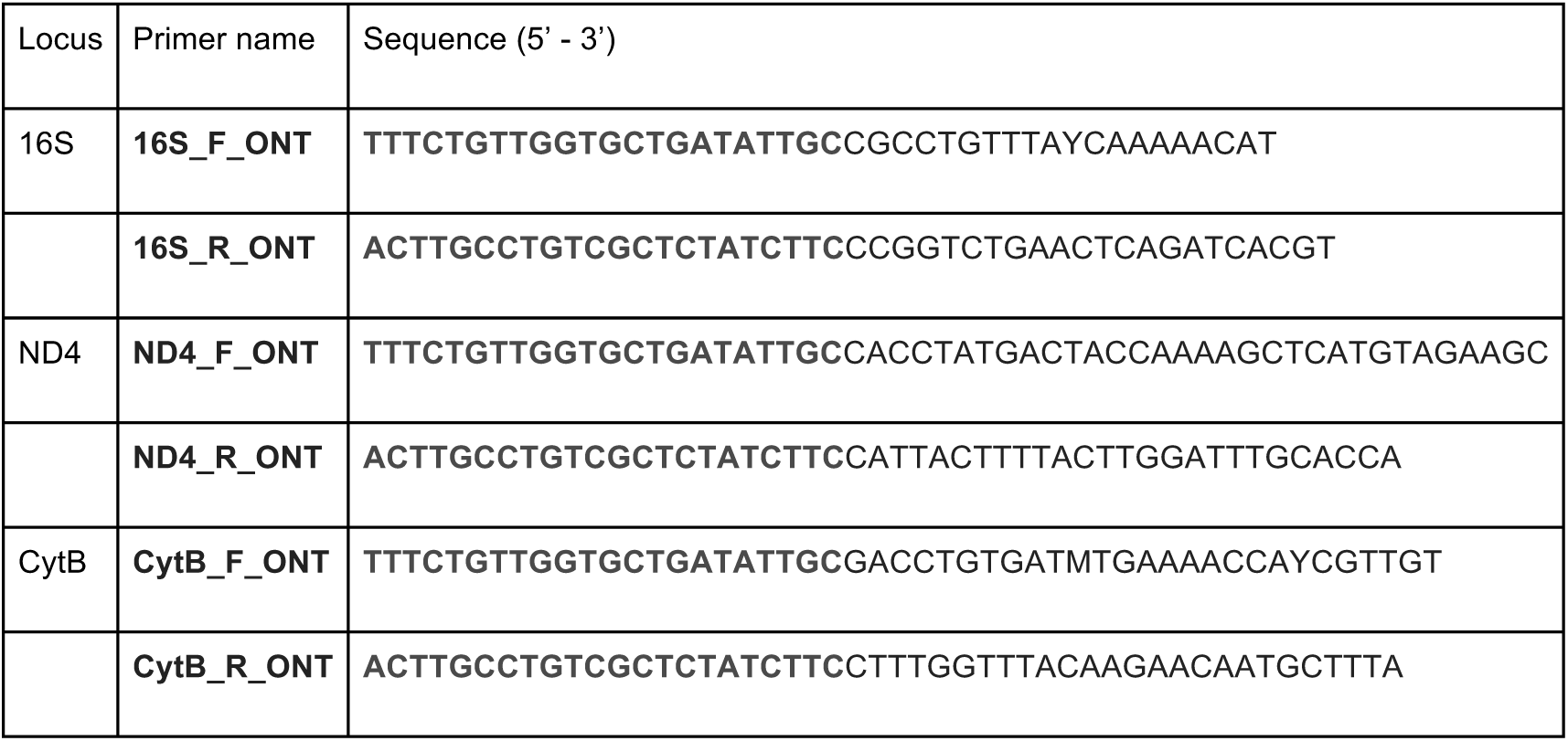
Primers used in this study. Bold nucleotides indicate the universal tailed ONT sequences for the barcoding kit.

**Supplementary Table 3.** Subsampling report: impact of coverage on the consensus accuracy by randomly subsampling.

Subsampling scheme for viper
|  | Reads | Assembled reads | Mapped reads | Mismatches | Accuracy |
| --- | --- | --- | --- | --- | --- |
| **Set 1** | 30 | 25 | 30 | 1 deletion poly T (5) | 99.8% |
|  | 30 | 29 | 30 | 1 deletion poly T (6) | 99.8% |
|  | 30 | 26 | 30 | 3 deletions poly T (5, 6, 5) | 99.4% |
| **Set 2** | 100 | 57 | 100 | 1 deletion poly T (5) | 99.8% |
|  | 100 | 53 | 100 | 2 deletions poly T (5, 6) | 99.6% |
|  | 100 | 62 | 100 | - | **100%** |
| **Set 3** | 300 | 60 | 300 | 1 deletion poly T (5) | 99.8% |
|  | 300 | 62 | 300 | 1 deletion poly T (5) | 99.8% |
|  | 300 | 53 | 300 | 1 deletion poly T (5) | 99.8% |
| **Set 4** | 1000 | 29 | 1000 | 1 deletion poly T (5) | 99.8% |
|  | 1000 | 56 | 1000 | 1 deletion poly T (5) | 99.8% |
|  | 1000 | 9 | 1000 | - | **100%** |

Subsampling scheme for gecko 1
|  | Reads | Assembled reads | Mapped reads | Mismatches | Accuracy |
| --- | --- | --- | --- | --- | --- |
| **Set 1** | 30 | 30 | 30 | 3 deletions poly A/A/C (5/4/5) | 99.4% |
|  | 30 | 17 | 30 | 2 deletions poly A/C (5/5) | 99.6% |
|  | 30 | 15 | 30 | 1 insertion poly A (5), 1 deletions poly C (5) | 99.8% |
| **Set 2** | 100 | 62 | 100 | one T instead of C (2Cs) | 99.8% |
|  | 100 | 26 | 100 | - | **100%** |
|  | 100 | 51 | 100 | - | **100%** |
| **Set 3** | 300 | 52 | 299 | - | **100%** |
|  | 300 | 56 | 300 | - | **100%** |
|  | 300 | 7 | 299 | - | **100%** |
| **Set 4** | 1000 | 1 | 999 | 1 C instead of T (CT) | 99.8% |
|  | 1000 | 57 | 999 | - | **100%** |
|  | 1000 | 57 | 998 | - | **100%** |

## Supplementary Material - Bioinformatics Commands

### Sequencing and base calling

The library was sequenced on the MinION platform using an offline MinKnow version without local base calling. At the time our team departed for the field only a Windows MinKnow offline version was available, thus we carried out sequencing and base calling on a Windows laptop. The analysis was subsequently performed using a Linux Ubuntu system. The sequencing was carried out using a MinION R9.5 flow cell. We then used Albacore 1.2.5 for local base calling and demultiplexing.

> -i E:\2017_Ecudador_Barcode_7_12_17 - t 2 - s E:\2017_Ecudador_Barcode_7_12_17 - f

FLO-MIN107 — kit SQK-LSK108 — recursive — barcoding — output_format fast5 In order to be able to perform sequence polishing we outputted fast5 in Albacore. The fast5 files for each individual barcode were subsequently converted to fastq files using Nanopolish.

> nanopolish extract - b albacore — fastq barcode01/ > barcode01_eyelash_palm-pitviper_16S.fastq

Nanopolish also adds a directory path to the fastq header in order to use the fast5 information later in the polishing.

Next we employed different consensus sequence generation methods, namely reference-based mapping, reference-based mapping using an ONT read as reference and de novo assembly of the individual amplicon sequences.

### Read Filtering

> cat barcode01_eyelash_palm-pitviper_16S.fastq | NanoFilt - q 15 - l 500 > barcode01_eyelash_palm-pitviper_16S_filtered.fastq

### De-novo assembly of the Amplicons (Preferred Option)

W created the consensus sequence *de novo*, so without the use of a reference sequence. Here we used the genome assembler Canu with adapted parameters for shorter reads. *De-novo* assembly of the amplicons is preferred over reference-based mapping (see below), since it does not introduce any biases by mapping it to a reference, which might have indels compared to the reads.

To perform a *de novo* assembly for each amplicon using canu we run:

> canu - p barcode01_eyelash_palm-pitviper_16S_canu - d barcode01_eyelash_palm-pitviper_16S_canu genomeSize=1000 minReadLength=100 minOverlapLength=50 - nanopore-raw barcode01_eyelash_palm-pitviper_16S.fastq
>
> tgStoreDump - T unitigging/barcode01_eyelash_palm-pitviper_16S_canu.ctgStore 2 - G unitigging/ barcode01_eyelash_palm-pitviper_16S_canu.gkpStore-consensus-fasta-tig 1 > barcode01_eyelash_palm-pitviper_16S_canu.bestcontig.fasta

Alternatively, we saved all contigs to a file using:

> tgStoreDump - T unitigging/barcode01_eyelash_palm-pitviper_16S_canu.ctgStore 2 - G unitigging/ barcode01_eyelash_palm-pitviper_16S_canu.gkpStore - consensus - fasta > barcode01_eyelash_palm-pitviper_16S_canu.contigs.fasta

We then created consensus sequences the same commands as described below (bwa index, bwa mem, samtools view, samtools sort, samtools index and nanopolish variants). We used the first contig from canu as the reference for the mapping. We further performed mapping against a reference that contained all canu generated contigs to check how many reads map to the different contigs. The first contig always showed the highest number of reads.

In the last step we removed adapters and the priming sites using cutadapt.

> cutadapt - g CGCCTGTTTAYCAAAAACAT…ACGTGATCTGAGTTCAGACCGG - o pitviper_filtered_canu.best.contig_cut.fasta
>
> pitviper_filtered_canu.best.contig.fasta

### Standard Reference-based Mapping

For Option 2 we mapped the reads to a reference sequence downloaded from NCBI (here GenBank Accession KC847257). In the first step, we indexed the reference.

bwa index KC847257_bothriechis_schlegelii.fasta We then mapped the reads onto the indexed reference using BWA mem. The mem algorithm was specifically designed for mapping reads to a divergent reference. Here we used bwa 0.7.12, which provides an option - d ont2d, which employs the following filters-k14-W20-r10-A1-B1-O1-E1-L0 in order to optimize the mapping for Oxford Nanopore read data.

> bwa mem - x ont2d KC847257_bothriechis_schlegelii.fasta barcode01_eyelash_palm-pitviper_16S.fastq > barcode01_eyelash_palm-pitviper_16S.sam

Here KC847257_bothriechis_schlegelii.fasta was the reference that we used in the mapping and barcode01_eyelash_palm-pitviper_16S.fastq the read data. In the next step we used samtools 1.3.1 to convert the mapping in sam format to a bam format, which performs block compression to reduce the file size. Mapping files in bam format are standardly used in downstream analysis.

> samtools view - bS barcode01_eyelash_palm-pitviper_16S.sam > barcode01_eyelash_palm-pitviper_16S.bam

We then sorted and indexed the bam file using samtools to allow rapid random access for indexing queries.

> samtools sort barcode01_eyelash_palm-pitviper_16S.bam - o barcode01_eyelash_palm-pitviper_16S.srt.bam samtools index barcode01_eyelash_palm-pitviper_16S.srt.bam

In the next step, we called the consensus sequence using ANGSD v. 0.911-51-g57d0264

> angsd - doFasta 3 - doCounts 1 barcode01_eyelash_palm-pitviper_16S.srt.bam - out barcode01_eyelash_palm-pitviper_16S_angsd

The - doFasta 3 option performs consensus calling using highest effective depth (EBD; Wang et al. 2013). The fasta file can be unzipped afterwards using “gzip - d file”.

In the next steps we mapped all the read data back onto the created consensus sequence to enable consensus polishing using nanopolish (same commands as before: bwa index, bwa mem, samtools view, samtools sort and samtools index). Finally we created a polished consensus using nanopolish.

> nanopolish variants — consensus=barcode01_eyelash_palm-pitviper_16S_cons.fasta — bam barcode01_eyelash_palm-pitviper_16S.srt.bam — genome barcode01_eyelash_palm-pitviper_16S_angsd.fasta — reads barcode01_eyelash_palm-pitviper_16S.fastq

Wang, Y., Lu, J., Yu, J., Gibbs, R. A., & Yu, F. (2013). An integrative variant analysis pipeline for accurate genotype/haplotype inference in population NGS data. *Genome Research*, *23*(5), 833-842. http://doi.org/10.1101/gr.146084.112

**Calculations for cost in this study, based on expenses as of July, 2017**:

- ONT starter kit: $1000
- ONT 12 barcode kit: $250 for 6 library run, therefore each library per run = $41.7
- 2 barcode libraries generated in this study = (41.7 x 2) + $1000 (ONT starter pack) = $1083.4
- Number of barcodes generated by 2 rounds of sequencing: 12 x 2 = 24 barcodes
- $1083.4 / 24 barcodes = $45.1 per barcode

